# HTSeq – A Python framework to work with high-throughput sequencing data

**DOI:** 10.1101/002824

**Authors:** Simon Anders, Paul Theodor Pyl, Wolfgang Huber

## Abstract

**Motivation:** A large choice of tools exists for many standard tasks in the analysis of high-throughput sequencing (HTS) data. However, once a project deviates from standard work flows, custom scripts are needed.

**Results:** We present HTSeq, a Python library to facilitate the rapid development of such scripts. HTSeq offers parsers for many common data formats in HTS projects, as well as classes to represent data such as genomic coordinates, sequences, sequencing reads, alignments, gene model information, variant calls, and provides data structures that allow for querying via genomic coordinates. We also present htseq-count, a tool developed with HTSeq that preprocesses RNA-Seq data for differential expression analysis by counting the overlap of reads with genes.

**Availability:** HTSeq is released as open-source software under the GNU General Public Licence and available from http://www-huber.embl.de/HTSeq or from the Python Package Index https://pypi.python.org/pypi/HTSeq.

## 1. INTRODUCTION

The rapid technological advance in high-throughput sequencing (HTS) has led to the development of many new kinds of assays, each of which requires the development of a suitable bioinformatical analysis pipeline. For the recurring “big tasks” in a typical pipeline, such as alignment and assembly, the bioinformatics practitioner can chose from a range of standard tools. For more specialised tasks, and in order to interface between existing tools, customised scripts often need to be written.

Here we present HTSeq, a Python library to facilitate the rapid development of scripts for processing and analysis of high-throughput sequencing (HTS) data. HTSeq includes parsers for common file formats for a variety of types of input data and is suitable as a general platform for a diverse range of tasks. A core component of HTSeq is a container class that simplifies working with data associated with genomic coordinates, i.e., values attributed to genomic positions (e.g., read coverage) or to genomic intervals (e.g., genomic features such as exons or genes). Two stand-alone applications developed with HTSeq are distributed with the package, namely htseq-qa for read quality assessment and htseq-count for preprocessing RNA-Seq alignments for differential expression calling.

Most of the features described in the following sections have been available since the initial release of the HTSeq package in 2010. Since then, the package and especially the htseq-count script have found considerable use in the research community. The present article provides a description of the package and also reports on recent improvements.

HTSeq comes with extensive documentation, including a tutorial that demonstrates the use of the core classes of HTSeq and discusses several important use cases in detail. The documentation, as well as HTSeq’s design, is geared towards allowing users with only moderate Python knowledge to create their own scripts, while shielding more advanced internals from the user.

## 2 COMPONENTS AND DESIGN OF HTSEQ

### 2.1 Parser and record objects

HTSeq provides parsers for reference sequences (FASTA), short reads (FASTQ), short-read alignments (the SAM/BAM format and some legacy formats), and for genomic feature, annotation and score data (GFF/GTF, VCF, BED, Wiggle).

Each parser is provided as a class which whose objects are tied to a file name or open file or stream and work as iterator generators, i.e., they may be used in the head of a *for* loop and will yield a sequence of record objects that are taken up by the loop variable. These record objects are instances of suitable classes to represent the data records. Where appropriate, different parsers will yield the same type of record objects. For example, the record class *SequenceWithQualities* is used whenever sequencing read with base-call qualities needs to be presented, and hence yielded by the *FastqParser* class and also present as a field in the *SAM Alignment* objects yielded by *SAM Reader* or *BAM Reader* parser objects (Figure 1). Specific classes (*GenomicPosition* and *GenomicInterval*) are used to represent genomic coordinates or intervals, and these are guaranteed to always follow a fixed convention (namely, following Python conventions, zero-based, with intervals being half-open) and parser classes take care to apply appropriate conversion when the input format uses different convention. The reverse is true for functions to write files.

**Fig. 1.**
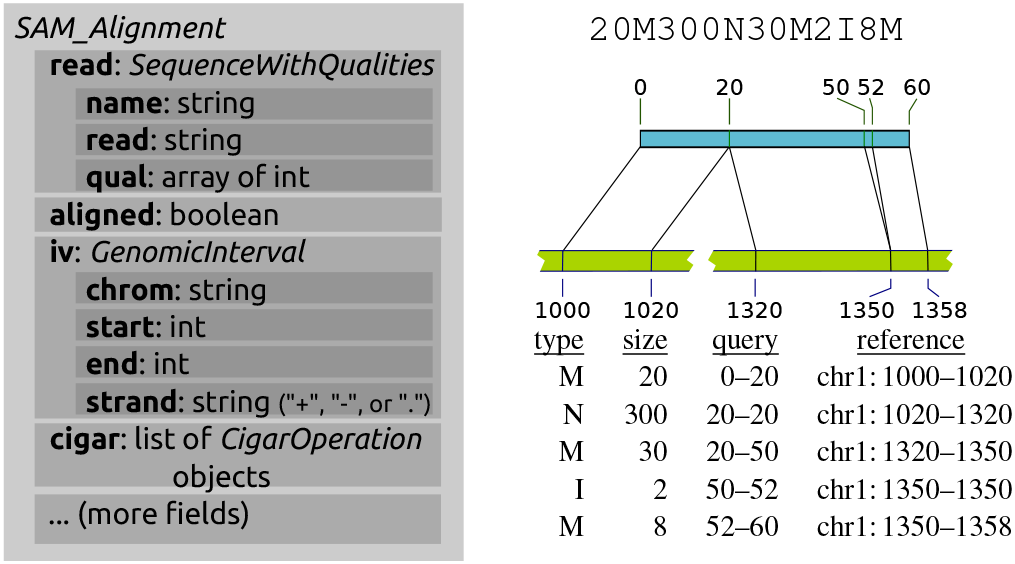
(a) The *SAM_Alignment* class as an example of an *HTSeq* data record: Subsets of the content are bundled in object-valued fields, using classes (here *SequenceWithQualities* and *GenomicInterval*) that are also used in other data records to provide a common view on diverse data types. (b) The *cigar* field in a *SAM_alignment* object presents the detailed structure of a read alignment as a list of *CigarOperation*. This allows for convenient downstrem processing of complicated alignment structures, such as the one given by the cigar string on top and illustrated in the middle. Five *CigarOperation* objects, with slots for the column of the table (bottom) provide the data from the cigar string, along with the inferred coordinates of the affected regions in read (“query”) and reference.

To offer good performance, large parts of *HTSeq*, are written in *Cython* (Behnel *et al.*, 2011), a tool to translate Python code augmented with type information to *C*. While the code for reading and writing all text-based formats, including text SAM files, is written in Python/Cython and hence has no external dependencies, the classes *BAM_Reader* and *BAM_Writer* wrap around functionality from *PySam* (Heger *et al.*) and are available only if that package has been installed.

The *SAM_Alignment* class offers functionality to facilitate dealing with complex, e.g. gapped, alignments (Figure 1b), with multiple aligments and with paired-end data. The latter is challenging because, in the SAM format, an alignment of a read pair is described by a pair of alignment records, which cannot be expected to be adjacent to each other. HTSeq provides a function, *pair_SAM_alignments_with_buffer*, to pair up these records by keeping a buffer of reads whose mate has not yet been found, and so facilitates processing data on the level of sequenced fragments rather than reads.

## 2.2 The GenomicArray class

Data in genomics analyses is often associated with positions on a genome, i.e., coordinates in a reference assembly. One example for such data is read coverage: for each base pair or nucleotide of the reference genome, the number of read alignments overlapping that position are stored. Similarly, gene models and other genomic annotation data can be represented as objects describing features such as exons that are associated with genomic intervals, i.e., coordinate ranges in the reference.

A core component of HTSeq is the class *GenomicArray*, which is a container to store any kind of genomic-position dependent data. Conceptually, each base-pair position on the genome can be associated with a value that can be efficiently stored and retrieved given the position, where the value can be both a scalar type, such as a number, or a more complex Python object. In practice, however, such data is often piecewise constant, and hence, the class internally uses a tree structure to store “steps”, i.e., genomic intervals with a given value. This has been implemented in *C++*, building on the *map* template of the C++ standard library, which is typically realized as a red-black tree (Josuttis, 1999). To link C++ and Python code, we used SWIG (Beazley *et al.*, 1996). Alternatively, the class also offers a storage mode based on NumPy arrays (van der Walt *et al.*) to accommodate dense data without steps. If such data becomes to large to fit into memory, NumPy’s *memmap* feature may be used, which swaps currently unused parts of the data out to disk. The choice of storage back-end is transparent, i.e., if the user changes it, no changes need to be made in the code using the *GenomicArray* objects.

A sub-class of *GenomicArray*, the *GenomicArrayOfSets* is suitable to store objects associated with intervals that may overlap, such as genes or exons from a gene model reference. This is implemented using Python sets (Figure 2): Each step’s value is a set of references to the actual objects. When data is inserted into the array, steps gets split and sets get duplicated as needed. When querying an interval, the sets overlapped by the query interval are returned, and their union will contain all objects overlapped by the query interval.

**Fig. 2.**
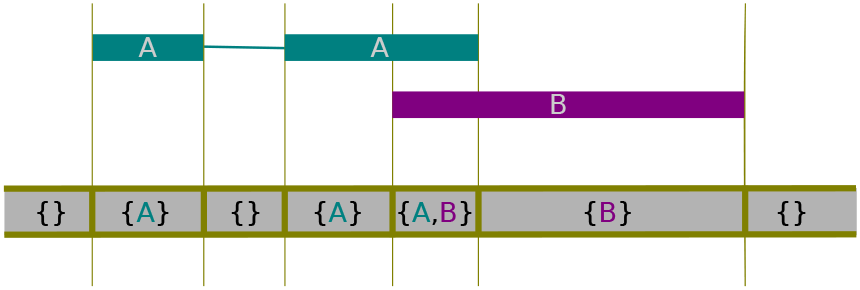
Using the class *GenomicArrayOfSets* to represent overlapping annotation metadata. The indicated features are assigned to the array, which then represents them internally as steps, each step having as value a set whose elements are references to the features overlapping the step.

## 3 DOCUMENTATION AND CODING STRATEGIES

HTSeq comes with extensive documentation to guide developers. Care has been taken to expect only moderate experience with Python from the reader. A “Tour” offers an overview over the classes and principles of HTSeq by demonstrating their use in simple examples. Then, two common use cases are discussed in detail to show how HTSeq can be applied to complex tasks.

The first use case is that of aggregate coverage profiles: Given ChiP-Seq data, e.g. from histone marks, we want to calculate the profile of these marks with respect to specific positions in the genome, such as transcription start sites (TSSs), by aligning coverage data in windows centred on the TSSs and averaging over the TSSs of all genes or a subset thereof. In this use case, one needs to integrate information from two position-specific data sources, namely a list of TSSs obtained from annotation data and the aligned reads. Hence, one may either iterate through the reads first, store this information in a *GenomicArray* and then use position-specific access to it when iterating through the list of TSSs, or, first store the TSSs in a *GenomicArray* and use this afterwards when iterating through the reads. In either case, one data set is kept in memory in a form allowing for fast random access, while the other is iterated through with only summary information being kept. These approaches are prototypical for scripts built on HTSeq and hence explained and demonstrated in detail in the documentation (Section “A detailed use case: TSS plots”).

The second use case discussed in detail is that of counting for each gene in a genome how many RNA-Seq reads map to it. In this context, the HTSeq class *CigarOperation* is demonstrated, which represents complex alignments in a convenient form (Figure 1b). This section of the documentation also explains HTSeq’s facilities to handle multiple alignments and paired-end data.

The remainder of the documentation provides references for all classes and functions provided by HTSeq, including those classes not used in the highlighted use cases of the tutorial part, such as the facilities to deal with variant-call format (*VCF*) files.

## 4 HTSEQ-COUNT

We distribute two stand-alone scripts with HTSeq, which can be used from the shell command line, without any Python knowledge, - and also illustrate potential applications of the HTSeq framework. The script *htseq-qa* is a simple tool for initial quality assessment of sequencing runs. It produces plots which summarise the nucleotide compositions of the positions in the read and the base-call qualities.

The script *htseq-count* is a tool for RNA-Seq data analysis: Given a SAM/BAM file and a GTF or GFF file with gene models, it counts for each gene how many aligned reads overlap its exons. These counts can then be used for gene-level differential expression analyses using methods such as *DESeq2* (Love *et al.*, 2014) or *edgeR* (Robinson *et al.*, 2010). As the script is designed specifically for *differential* expression analysis, only reads mapping unambiguously to a single gene are counted while reads aligned to multiple positions or overlapping with more than one gene are discarded. To see why this is desirable, consider two genes with some sequence similarity, one of which is differentially expressed while the other one is not. A read that maps to both genes equally well should be discarded, because if it were counted for both genes, the extra reads from the differentially expressed gene may cause the other gene to be wrongly called differentially expressed, too. Another design choice made with the downstream application of differential expression testing in mind is to count fragments, not reads, in case of paired-end data. This is because the two mates originating from the same fragment provide only evidence for one cDNA fragment and should hence be counted only once.

As the *htseq-count* script has found widespread use over the last three years, we note that we recently replaced it with an overhauled version, which now allows processing paired-end data without the need to sort the SAM/BAM file by read name first. See the documentation for a list of all changes to the original version.

## 5 DISCUSSION

HTSeq aims to fill the gap between performant but monolithic tools optimised for specialised tasks and the need to write data processing code for HTS application entirely from scratch. For a number of the smaller tasks covered by HTSeq, good stand-alone solutions exist, e.g. *FastQC* (Andrews *et al.*, 2011) for quality assessment or *Trimmomatic* (Bolger *et al.*, 2014) for trimming of reads. If the specific approaches chosen by the developers of these tools are suitable for a user’s application, they are easier to use. However, the need to write customised code will inevitably arise in many projects, and then, HTSeq aims to offer advantages over more narrow programming libraries that focus on specific file formats, e.g. *PySam* (Heger *et al.*) and *Picard* (Wysoker *et al.*) for SAM/BAM files, by integrating parsers for many common file formats and fixing conventions for data interchange between them. For R developers, similar functionality is now available within the *Bioconductor* project (Gentleman *et al.*, 2004) with packages like *Rsamtools* (Morgan *et al.*) and *GenomicRanges* (Lawrence *et al.*, 2013). Within Python, HTSeq complements Biopython (Cock *et al.*, 2009), which provides similar functionality for more “classic” bioinformatics tasks such as sequence analysis and phylogenetic analyses but offers little support for HTS tasks.

While most uses of HTSeq will be the development of custom scripts for one specific analysis task in one experiment, it can also be useful for writing more general tools. The *htseq-count* script, for example, prepares a count table for differential expression analysis, a seemingly easy task, which, however, becomes complicated when ambiguous cases have to be treated correctly. Despite being written in Python, *htseq-count* offers decent performance: Tested on a standard laptop computer, htseq-count (version 0.6.1) processed about 1.2 million reads (0.6M read pairs) per minute, using about 250 MB of RAM to hold the human gene annotation in memory. When the file was sorted by position rather than read-name, so that mate pairs were not in adjacent record, processing time increased to a bit less then twice as much, and, for a SAM file of 26 GB, less than 450 MB of additional space in RAM were needed for the buffer holding reads with outstanding mates.

When HTSeq was first released in 2010, *htseq-count* was the first comprehensive solution for this task, and has since then been widely used. Recently, further tools for this task have become available, including the *summarizeOverlap* function in the *GenomicRanges* Bioconductor package (Lawrence *et al.*, 2013) and the stand-alone tool *featureCount* (Liao *et al.*, 2014), which achieves very fast run times due to being implemented in C. In a recent benchmark, Fonseca *et al.* (2014) compared *htseq-count* with these other counting tools and judged the accuracy of *htseq-count* favourably. Nevertheless, neither *htseq-count* nor the other tools offer much flexibility to deal with special cases, which is why the HTSeq documentation (section “Counting reads”) discusses in detail how users can write their own scripts for this important use case.

Interval queries are a recurring task in HTS analysis problems, and several libraries now offer solutions for different programming languages, including BEDtools (Quinlan and Hall, 2010; Dale *et al.*, 2011) and IRanges/GenomicRanges (Lawrence *et al.*, 2013). Typically, these methods take two lists of intervals and report overlaps between them. HTSeq uses a different paradigm, namely that one list of intervals is read in and stored in a *GenomicArrayOfSets* object, and then the other intervals are queried one by one, in a loop. This explicit looping can be more intuitive; one example is the read counting problem discussed above, where split reads, gapped alignments, ambiguous mappings etc. cause much need for treatment of special cases that is addressed by branching statements within the inner loop.

In conclusion, HTSeq offers a comprehensive solution to facilitate a wide range of programming tasks in the context of high-throughput sequencing data analysis.

*Funding:* SA and WH acknowledge support from the European Union via the FP6 network *Chromatin Plasticity* and the FP7 project *Radiant*.

